# Complex patterns of sex-biased demography in canines

**DOI:** 10.1101/362731

**Authors:** Tanya N. Phung, Robert K. Wayne, Melissa A. Wilson Sayres, Kirk E. Lohmueller

## INTRODUCTION

Studies of genetic variation have shown that the demographic history of dogs has been extremely complex, involving multiple bottleneck and admixture events. However, existing studies have not explored the variance in the number of reproducing males and females, and whether it has changed across evolutionary time. While male-biased mating practices, such as male-biased migration and multiple paternity, have been observed in wolves, recent breeding practices could have led to female-biased mating patterns in breed dogs. In addition, breed dogs are thought to have experienced the popular sire effect, where a small number of males father many offspring with a large number of females. Here we use genetic variation data to test how widespread sex-biased mating practices in canines are during different time points throughout their evolutionary history. Using whole genome sequence data from 33 dogs and wolves, we show that patterns of diversity on the X chromosome and autosomes are consistent with a higher number of reproducing males than females over ancient evolutionary history in both dogs and wolves, suggesting that mating practices did not change during early dog domestication. In contrast, since breed formation, we found evidence for a larger number of reproducing females than males in breed dogs, consistent with the popular sire effect. Our results confirm that the demographic history of canines has been complex, with unique and opposite sex-biased processes occurring at different times. The signatures observed in the genetic data are consistent with documented sex-biased mating practices in both the wild and domesticated populations, suggesting that these mating practices are pervasive.

Dogs were the first animals known to be domesticated and have lived alongside humans and shared our environment ever since^1^. There is tremendous interest in understanding their genetics and evolutionary history^2–5^. Many studies have shown that dogs have a complex evolutionary history; they experienced a population size reduction (i.e. bottleneck) associated with domestication and additional breed-specific bottlenecks associated with breed formation during the Victorian era^6^. In addition to bottleneck events, dogs experienced admixture with wolves during the domestication process^7^. Studies have disagreed about the process of domestication, including when, where, and how many times dogs were domesticated^2,8–12^. However, despite the extensive work on understanding dog demographic history, existing studies have not explored the population history of males and females across dog domestication. Departures from an equal number of reproducing males and females are called sex-biased demographic processes, and leave signatures in the genome (reviewed in Wilson Sayres 2018^13^). Previous ecological and field studies suggested that mating practices have been sex-biased in canines. In the wild populations, vonHoldt et al. (2008) observed that in some cases, Yellowstone male wolves would migrate to an existing wolf pack to mate with the alpha female when the alpha male dies^14^. The migration into an existing wolf pack is therefore male-biased. An additional source of male biased migration may come from male wolves called “Casanova wolves”. These wolves leave their natal packs and visit a nearby wolf pack around mating season to mate with the subordinate females^15^. Lastly, there has also been evidence of multiple paternity in Ethiopian wolves and foxes^16,17^. In the domesticated populations, it is thought that more females contributed to breed formation than males, indicating female-biased processes^18^. In addition, recent reproductive practices, such as the popular sire effect, which involves a small number of males reproducing with a large number of females can lead to female-biased demography^19^. Despite these observations of mating practices suggesting the numbers of reproducing males and females has been unequal during canid evolution, it is unclear how pervasive these processes are, and which have had the dominant effect on shaping patterns of diversity.

To test how widespread sex biased demography has been throughout canid evolution, we calculated and compared measures of genetic diversity on the X chromosome to those on the autosomes. This ratio has been termed Q in Emery et al. (2010) and we will use this notation throughout^20^. In male-heterogametic sex-determining systems (XX/XY) with equal numbers of reproducing males and females, there are three copies of the X chromosome for every four copies of the autosomal genome. Therefore, in a constant size population without any natural selection or sex-biased processes,Q is expected to be 0.75 (reviewed in Webster and Wilson Sayres 2016^21^). Specifically, Q = N_X_/ N_A_ ≅ 0.75. Deviations from this expected ratio could be indicative of sex-biased processes. If Q < 0.75, there are fewer copies of the X chromosome than expected, suggesting a larger number of reproducing males than reproducing females, indicative of male-biased processes. If Q > 0.75, there are more copies of the X chromosome than expected, suggesting a larger number of reproducing females than reproducing males, indicative of female-biased processes.

Studies comparing measures of genetic diversity between the X chromosome and autosomes have resulted in many insights into the evolutionary history of humans. Hammer et al. (2008) computed Q by fitting a model of demographic history to the ratio in the mean of genetic diversity within the X chromosome and autosomes: 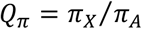^22^. They found that Q_π_ is greater than 0.75 in all human populations examined, suggesting female-biased processes that have led to more reproducing females than males during human evolutionary history^22^. Later, Keinan et al. (2009)^23^ computed Q by calculating the ratio in fixation index, F_ST_, between the X chromosome and the autosomes: 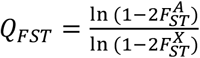. They found that Q_FST_ is less than 0.75 only when comparing a non-African population to an African population^23^. This result suggests that there was a male-biased migration out of Africa, where there were more reproducing males than females. Even though these two studies came to different conclusions regarding the sex ratio in human history, a later study reconciled these seemingly disparate findings by demonstrating that Q can detect bias in sex ratios at different timescales, depending on whether it is calculated from genetic diversity (Q_π_) or the fixation index (Q_FST_)^20^. Specifically, Q_π_ can detect sex bias in ancient timescales, which is before or immediately after the split between populations, whereas Q_FST_ detects sex-biased demography on recent timescales, after the populations split from each other^20^. Emery et al. (2010) reconciled results from Hammer et al. (2008) and Keinan et al. (2009) by showing that evolutionary processes within human history are consistent with an earlier female bias followed by a male bias during the migration of some humans out of Africa^20^. Additionally, direct comparisons of the two studies were complicated by linked selection on the X chromosome^24,25^. In addition to humans, comparing the genetic diversity between the X chromosome and autosomes has also been used to study sex-biased processes in many other species^13^.

Given how examining patterns of genetic diversity on the X chromosome and the autosomes has facilitated our understanding of sex-biased demography in other species and what has been observed regarding sex-biased mating practices in canines, we wanted to test how widespread these mating practices are throughout different time points during canine evolutionary history. We utilized whole-genome sequences of 21 dogs and 12 wolves. Using the estimator of the effective sex ratio based on nucleotide diversity, we found that Q_π_ is less than 0.75 in both dogs and wolves, indicative of an ancient male bias either in the shared ancestral population, or immediately after their split. We then inferred the effective sex ratio in a population genetic model, demonstrating that a population size reduction by itself cannot generate the empirical patterns. Rather, a male-biased sex ratio was needed in conjunction with a population size reduction to recapitulate empirical patterns. Finally, using the estimator of the effective sex ratio based on the fixation index, we showed that while the demographic history in wolves has remained male-biased in recent history, the demographic history in dogs has changed from male-biased in the ancient timescale to female-biased in recent times. These results add to our current understanding about the canine demographic history and suggest the need to incorporate sex-biased demography in future studies.

## RESULTS

### Description of the data

We collected a dataset of 33 female canid whole genomes that include 4 German Shepherds, 5 Tibetan Mastiffs, 12 dog individuals from a variety of breeds, 6 Arctic Wolves, and 6 Grey Wolves (Supplementary Table 1). The German Shepherd and Tibetan Mastiff data were sequenced by Gou et al. (2014)^26^ and the *fastq* files were downloaded from NCBI SRA. We combined 12 high coverage (>15X) whole genome sequences of female dogs from multiple breeds that were included in Marsden et al. (2016)^27^ because we were interested in how results differ between using a group of one breed versus using a group consisting of multiple breeds. We named this pooled group the “Pooled Breed Dogs”. The Arctic Wolf data were sequenced by Robinson et al (Submitted). These Arctic Wolves were located in Northern Canada (north of the Arctic circle). The longitudinal and latitudinal locations for these Arctic Wolves are included in Supplementary Table 1. We also used high coverage (>15X) whole genome sequences of female Grey Wolves from Marsden et al. (2016)^27^. Since these Grey Wolves originated from Europe, Asia, and Yellowstone, we named this population the “Pooled Grey Wolves”. Details about coverage and accession numbers for the individuals in this study are summarized in Supplementary Table 1.

### Estimating the effective sex ratio based on genetic diversity

Previous work has shown that dogs experience male mutation bias, where the mutation rate is higher in males compared to females due to more germline cell divisions in males at reproduction^28–30^. Male mutation bias has a significant impact on measurements of genetic diversity because it can inflate raw metrics of genetic diversity on the autosomes compared to on the X chromosome (reviewed in Webster and Wilson Sayres 2016^21^). To confirm that male mutation bias exists in our data, we computed male mutation bias for each population using dog-cat divergence (see Methods). We observed that the level of male mutation bias is around 2, which is consistent with previous reports^29,30^ (Supplementary Table 2). Therefore, we controlled for male mutation bias in all estimates of genetic variation by normalizing autosomal and X chromosome diversity by dog-cat divergence in the corresponding regions.

Natural selection is thought to be more efficient at reducing genetic diversity on the X chromosome than on the autosomes because males have only one X chromosome which is exposed directly to selection (reviewed in Webster and Wilson Sayres 2016^21^). To control for natural selection affecting the X chromosome more than the autosomes, we used regions of the genome in which mutations would be putatively neutral by removing sites that are functional. Specifically, we removed genic and conserved sites (see Methods).

To understand whether any evolutionary process has been sex-biased over ancient timescales, we computed Q_π_. We found that in both dog and wolf populations, Q_π_ is significantly less than 0.75 (Figure 1, No cM cutoff), suggesting a male-biased sex ratio, with more males reproducing relative to females.

**Figure 1:**
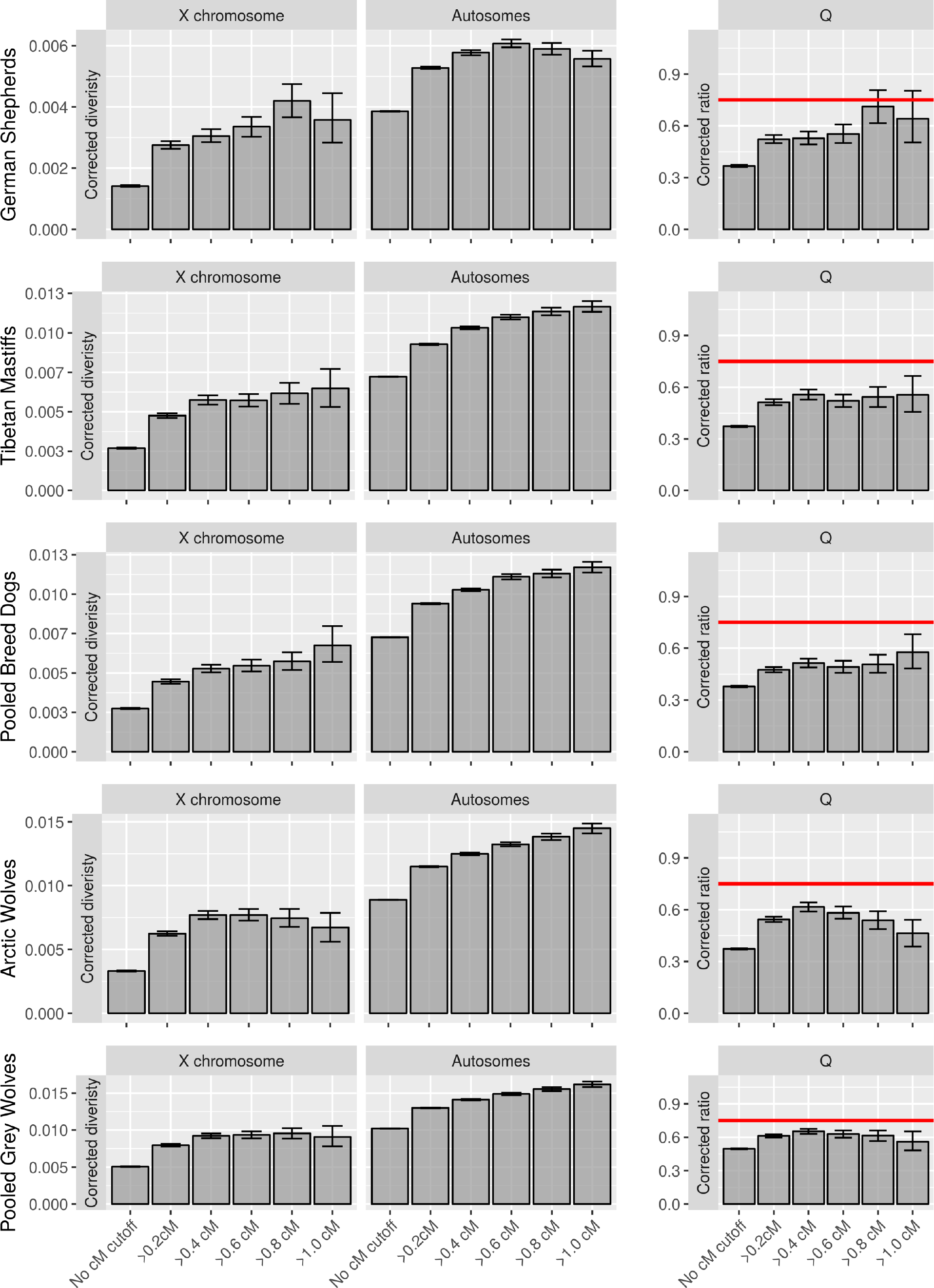
X-linked and autosomal genetic diversity across canids. Genetic diversity measured as the average pairwise differences between sequences (*π*) corrected for mutation rate variation using divergence (see Methods) on the X chromosome and autosomes in multiple canid populations. Q denotes the ratio of *π* on the X chromosome to that of the autosomes. The horizontal red line denotes the null expectation of 0.75. Bins along the x-axis denote different filtering based on genetic distances from genes. Error bars denote 95% confidence intervals obtained through bootstrapping (see Methods).

Q_π_ of less than 0.75 could occur due to the effect of natural selection on linked neutral sites. Specifically, natural selection could have reduced diversity in linked neutral regions on the X chromosome more than on the autosomes, as seen in humans^23–25^. Further, it is possible that there is more constraint on noncoding regions near genes on the X chromosome than on the autosomes^31^. To measure how neutral diversity is affected by linked selection, we compared diversity on the X chromosome and autosomes in regions near genes versus putatively unconstrained regions 0.4 cM away from the nearest gene. Diversity increased more with increasing distance from genes on the X chromosome than on the autosomes, consistent with natural selection reducing diversity more on the X chromosome than on the autosomes near genes (Supplementary Table 3).

To test whether stronger linked selection acting on the X chromosome relative to the autosomes could cause Q_π_ to be less than 0.75, we expanded our filtering criteria to remove sites that are near genes, defined by genetic distance (see Methods). Since we did not know *a priori* what the minimum genetic distance would be required to obtain sites that are not affected by selection, we included several thresholds. We removed sites whose genetic distance to the nearest genes is less than 0.2 cM, 0.4 cM, 0.6 cM, 0.8 cM, and 1 cM. We observed that even after removing sites whose genetic distance to the nearest genes are less than 1 cM, Q_π_ is still less than the expected 0.75 in both dog and wolf populations, except for the German Shepherd (Figure 1). In the German Shepherd, when using the thresholds of 0.8 cM and 1 Q_π_ approaches 0.75. However, since there are significantly fewer sites and variants left after removing sites whose genetic distance to the nearest genes is less than 0.8 cM or 1 cM, we could not exclude the possibility that we are underpowered to detect any signal in the data (Supplementary Table 4). Nonetheless, these results suggest that while linked selection may partially account for Q_π_ of less than 0.75, especially in the German Shepherd, linked selection by itself cannot explain why Q_π_ is less than 0.75 across all dog and wolf populations. In sum, our results suggest that there has been male-biased sex ratios in both dogs and wolves over ancient evolutionary timescales.

### Inference of sex-biased demographic processes under population genetic models

Pool and Nielsen (2007) demonstrated that a Q_π_ of less than 0.75 could be explained by a reduction in population size even with an equal number of breeding males and females^32^. To test whether population bottlenecks can explain the reduction in diversity on the X chromosome, we fitted a demographic model that includes a bottleneck using the autosomal site frequency spectrum (SFS) (Supplementary Figure 1) and asked whether the best fitting demographic model on the autosomes could also account for the level of diversity on the X chromosome when using an N_X_/N_A_ ratio of 0.75. If a demographic model including a bottleneck by itself can generate a Q_π_ of less than 0.75, we would expect that scaling the population size of the X chromosome to be three-quarters that of the autosomes should result in a Q_π_ comparable to the empirical data. Additionally, we then employed a composite likelihood framework to directly infer the N_X_/ N_A_ ratio from the SFS while accounting for the complex non-equilibrium demography.

First, we fitted a demographic model that includes a bottleneck using the SFS on the autosomes using *fastsimcoal2*^33^ for each population considering regions of greater than 0.4 cM, 0.6 cM, 0.8 cM, and 1 cM from genes. We reasoned that we would not be able to exclude the role of selection when not removing sites near genes or using too small of a threshold (i.e. 0.2 cM). We also corrected for male mutation bias using mutation rates that we inferred from dog-cat divergence in the same windows (see Methods; Supplementary Table 2). The inferred demographic parameters that resulted in the best likelihood of the data are presented in Supplementary Table 5. To test whether the inferred demographic parameters can recapitulate the autosomal data, we used *fastsincoal2* to generate the expected SFSs. In all populations except the German Shepherds, across all thresholds examined, we observed that the SFSs generated using the inferred demographic parameters visually match with the empirical autosomal SFSs (Supplementary Figure 2). The differences in log-likelihood between the simulated SFSs and the empirical SFSs are also small (Supplementary Table 6), confirming our visual inspection of the fit of the demographic models. In addition, autosomal genetic diversity (π) computed from the demographic model is comparable to the empirical estimates of π(Supplementary Figure 3). Thus, these lines of evidence demonstrate that the inferred demographic parameters can recapitulate the empirical data on the autosomes, except for the more stringent filtering on the German Shepherd (See Supplementary Note 1).

To understand whether the demographic model including a bottleneck that was fitted to the autosomal data could account for the level of diversity on the X chromosome, we used the inferred demographic parameters to simulate the SFSs for the X chromosome. To account for the differences in population size between the X chromosome and the autosomes, we adjusted the population size on the X chromosome by a constant value which we called C, where N_X_ = CN_A_. If a bottleneck by itself without any sex biased demography can generate a Q_π_ of less than 0.75, we expected that using a @ value of 0.75 would recapitulate the empirical data. If a bottleneck model by itself is not sufficient to generate a Q_π_ of less than 0.75, and sex-biased processes need to be invoked, we expected that rescaling the population size on the X chromosome to be three-quarters of the population size on the autosomes would not fit well. Rather, a different value of C would yield a better fit.

To assess whether a null C value of 0.75 or a different C value yielded a better fit to the empirical SFSs on the X chromosome, we searched over a grid of C values. We found the maximum likelihood value of C for each population and filtering threshold. To do this, for each C ona grid of C values, we first calculated the population size on the X chromosome, which is N_X_=CN_A_. We then used *fastsimcoal2* to simulate an SFS and assess the fit by comparing the Poisson log-likelihood to the SFS on the X chromosome (see Methods). For each population and for each threshold, we found a set of C values that maximizes the likelihood of the data (Figure 2, Table 1, Supplementary Table 7).

**Figure 2:**
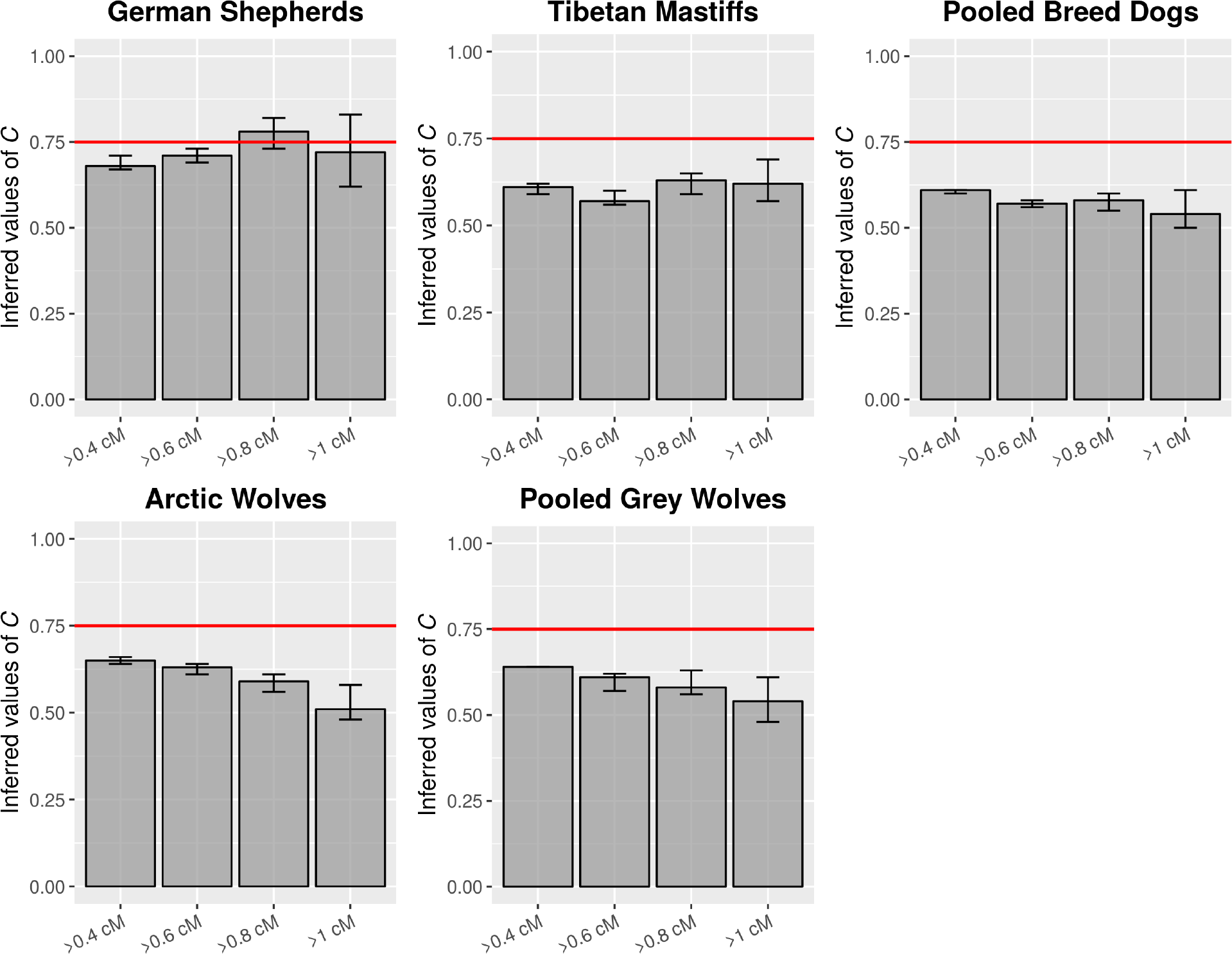
Effective population size estimates for multiple canid populations. Maximum likelihood estimates (MLEs) of the effective population size on the X chromosome relative to that of the autosomes (@ = N_X_/ N_A_) are shown for German Shepherds, Tibetan Mastiffs, Pooled Breed Dogs, Arctic Wolves and Pooled Grey Wolves with increasing distance from genes. Error bars denote approximate asymptotic 95% confidence intervals obtained as the parameter values within 2 log-likelihood unites of the MLE.

**Table 1.** Likelihood ratio tests comparing models of sex-biased demography in multiple canid populations. Likelihood ratio tests of the amount of sex-biased demography are shown when removing any sites whose genetic distance to the nearest genes is less than 0.4 cM. For the other thresholds, see Supplementary Table 7.

| Population | *C* | Log-likelihood | Likelihood ratio test | p-value |
| --- | --- | --- | --- | --- |
| German Shepherds | Null (*C* = 0.75) | 7460.957 | 34.779 | 3.69 X 10<sup>-9</sup> |
|  | Best (*C* = 0.68) | 7478.347 |  |  |
| Tibetan Mastiffs | Null (*C* = 0.75) | 16050.84 | 208.972 | 2.30 X 10<sup>-47</sup> |
|  | Best (*C* = 0.61) | 16155.32 |  |  |
| Pooled Breed Dogs | Null (*C* = 0.75) | 16875.26 | 249.627 | 3.13 X 10<sup>-56</sup> |
|  | Best (*C* = 0.61) | 17000.07 |  |  |
| Arctic Wolves | Null (*C* = 0.75) | 23035.46 | 133.341 | 7.62 X 10<sup>-31</sup> |
|  | Best (*C* = 0.65) | 23102.13 |  |  |
| Pooled Grey Wolves | Null (*C* = 0.75) | 30767.69 | 188.719 | 6.05 X 10<sup>-43</sup> |
|  | Best (*C* = 0.64) | 30862.05 |  |  |

With the exception of the German Shepherd at the most stringent filtering thresholds (>0.8 cM and >1 cM), we inferred that C is less than 0.75 for all population and filtering thresholds. When using a filtering threshold of 0.4 cM from genes, we found that C ranges from 0.61 to 0.68. The full model, where we inferred C for each comparison, fits the observed X chromosome SFS significantly better than a model where C is constrained to be 0.75 (Likelihood Ratio Tests > 30, p-value < 10^-8^; Table 1). Further, the null C value of 0.75 does not visually fit the SFSs on the X chromosome (Supplementary Figure 4, blue bars), suggesting that we can reject an equal number of reproducing males and females. Third, we observed that diversity on the X chromosome from simulating with a null C value of 0.75 overestimated the empirical X chromosome diversity (Supplementary Figure 5, blue bars). These results suggest that a model including both a bottleneck and a male-bias sex ratio can generate Q_π_ of less than 0.75 and recapitulate the observed SFSs and genetic diversity. Only in the German Shepherd population when using the most stringent threshold (>0.8 cM and >1 cM), can a demographic history including a bottleneck by itself generate a Q_π_ of less than 0.75.

### Female-biased sex ratio within dogs in recent history

Since estimates of sex ratios from levels of genetic diversity are sensitive to ancient sex-biased processes (prior to or immediately after the split between two species), we wanted to determine whether the pattern of male-biased contributions remained constant throughout the evolutionary history of canines^20^. To study sex-biased demography on recent timescales, we computed Q_FST_ for each pair of populations (see Methods). In the dog to dog comparison, we computed Q_FST_ between German Shepherds and Tibetan Mastiffs, between German Shepherds and Pooled Breed Dogs, and between Tibetan Mastiffs and Pooled Breed Dogs. We observed that Q_FST_ is greater than 0.75 for all three pairs and across all thresholds, suggesting a female-biased sex ratio within the dog populations in recent history (Figure 3 and Supplementary Figure 6). This is consistent with fewer reproducing males than females in the population since the formation of different dog breeds. In the wolf to wolf comparison, we computed Q_FST_ between Arctic Wolves and Pooled Grey Wolves. In contrast to the breed dogs, we found that Q_FST_ is less than 0.75 when using the thresholds of >0.4 cM and >0.6 cM, suggesting that a male-biased sex ratio has been maintained within the wolf populations in recent history (Figure 3 and Supplementary Figure 6). However, we noted that when using a more stringent threshold (>0.8 cM or >1 cM), Q_FST_ within wolves approaches 0.75 or greater than 0.75 (Supplementary Figure 6). We could not exclude the possibility that we are unable to detect a true signal in the data due to significantly fewer sites and variants left after the more stringent filtering (Supplementary Table 4). Overall, these results indicate that while the process within wolves has probably maintained a male-bias from ancient to recent history, the process within dogs has changed to female-bias, potentially because of breeding practices that have led to female-biased processes such as the popular sire effect.

**Figure 3:**
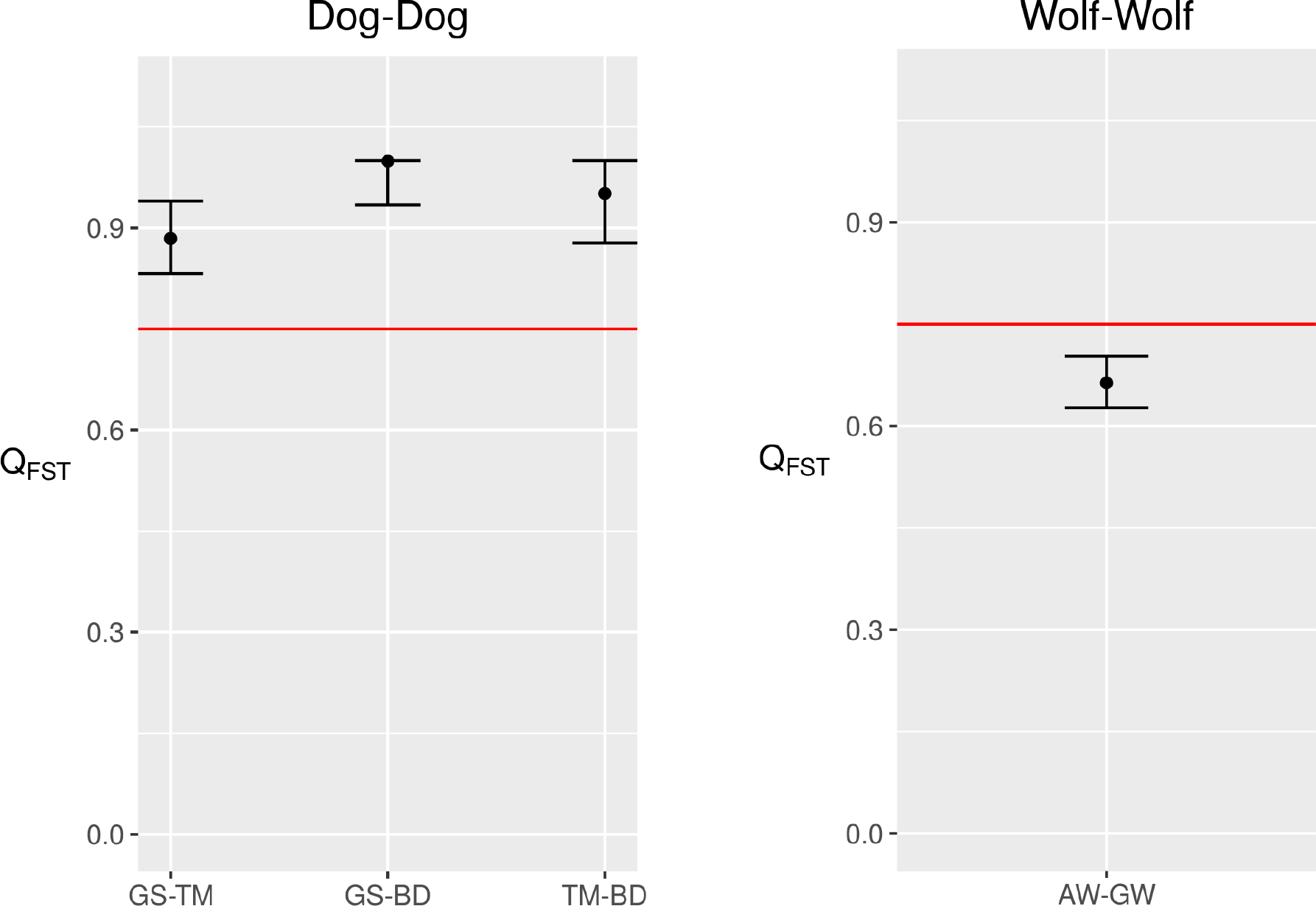
Sex biased demography on recent time scales. Estimates of the sex ratio for a pair of populations computed using *F_ST_* (see Methods) using a threshold of >0.6 cM to remove linked neutral sites are shown. The horizontal red line denotes the null expectation of 0.75. Error bars denote 95% confidence intervals obtained through bootstrapping (see Methods). Abbreviations: GS (German Shepherds), TM (Tibetan Mastiffs), BD (Pooled Breed Dogs), AW (Arctic Wolves), GW (Pooled Grey Wolf).

## DISCUSSION

In this study, we used two different statistics to estimate the ratio of reproducing males to females in canines and found that the demographic history of dogs and wolves has been sex-biased, but not always in the same direction. Estimating the sex ratio based on the levels of genetic diversity (Q_π_) from the X chromosome and autosomes showed a male-biased sex ratio in both dogs and wolves on an ancient timescale, which cannot be explained by linked selection or a population size reduction on its own (Figure 1 and Figure 2). Instead, in both dogs and wolves, there has been a larger number of reproducing males than females. In wolf packs, the alpha male and female are the dominant reproducers, but subdominant reproduction is common and may involve multiple fathers for a single litter^14^. Multiple paternity is a unique aspect of canid reproduction and may help drive a male bias in reproduction, as offspring of a single litter can only have a one mother, but may have multiple fathers and litter size may be as large as 16 individuals^34^. In addition, wolves migrating to existing wolf packs are predominantly male-biased^14^. Further, “Casanova wolves” who stay near a wolf pack during mating season to mate with the non-alpha females could also cause male-biased mating patterns^15^. Multiple paternity and male-biased migration likely occurred in early dogs, but under more recent controlled breeding, valuable sires would be the only father of a litter. Hence the controlled nature of breeding in modern dog breeds, and the focus on a subset of “popular” sires could drive the female bias in reproduction. The population sire effect also reduces the effective size of breeds and effects such as inbreeding further skew evolution in modern breeds.

In addition, we observed that determining the amount of bias based on the absolute value of Q_π_ by itself can lead to overestimation, because the reduction of diversity on the X chromosome due to a population size reduction is not accounted for. For example, in Tibetan Mastiff, when using a threshold of 0.6 cM to remove linked neutral sites, a Q_π_ of 0.52 suggests an N_X_ / N_A_ ratio of 0.52. However, we inferred a *C* value of 0.57 (confidence interval: 0.56-0.6) using our modelling framework, indicating that the sex ratio is higher than when just examining the absolute value of Q_π_. This difference exists because the estimate Q_π_ could be affected by a population size reduction differentially influencing diversity on the X and autosomes^32^, but our inference framework accounts for this effect. Our findings suggest that inferring the sex ratio in a model-based framework should yield a more accurate estimate than the absolute Q_π_^22^.

Our results add to the growing literature on the complex demographic history of dogs (reviewed in Freedman et al. 2016^3^ and Ostrander et al. 2017^4^). In addition to multiple episodes of bottleneck and admixture events, we now present evidence for sex-biased demographic processes. Furthermore, we provide evidence that sex-biased processes within dogs have changed throughout evolution, switching from a male-bias in ancient timescales to a female-bias in recent timescales, reflecting how modern breeding practices influence the sex ratio. To the best of our knowledge, this is the first genomic study of sex-biased demography in dogs. Some limitations in this study provide avenues for future work. First, our study was limited by the availability of high coverage (>15X coverage) whole-genome sequences of female individuals at the time of analysis. Future studies could utilize more female individuals and a variety of populations to understand whether there are differences in sex-biased processes between breeds. Second, future work could extend our modelling framework by including more complex demographic scenarios such as migration events to better capture the autosomal data, especially the German Shepherds. Finally, future studies could examine whether processes such as admixture with wolves or introgression has been sex-biased.

## METHODS

### Whole-genome sequence processing

We followed Genome Analysis Toolkit’s (GATK) documentation for variant discovery best practices^35–37^. Scripts used for processing whole-genome sequencing for each of the following steps can be found at https://github.com/tnphung/NGS_pipeline.

#### Data pre-processing for variant calling

First, we converted all fastq files to raw unmapped reads using Picard FastqToSam^38^. Second, we marked Illumina adapters using Picard MarkIlluminaAdapters^38^. Third, we mapped to the reference dog genome (canFam3) using bwa-mem^39^. Fourth, we marked duplicates using Picard MarkDuplicates^38^. We then recalibrated base quality scores using GATK where we performed three rounds of recalibration to obtain analysis-ready reads in BAM file format.

#### Variant calling with GATK

We used GATK Haplotype caller for variant calling^35–37^. We first generated a gVCF file for each individual. We then performed joint-genotyping for all 33 individuals in our study.

#### Filtering to obtain high quality sites

To obtain sites that are high confidence, we retained sites whose depth (annotated as DP in VCF file format) is between 50% and 150% of the mean depth across all sites. In addition, we only kept sites that were genotyped in all 33 individuals (i.e. the total number of alleles in called genotypes, AN, is equal to 66).

#### Variant filtering

We obtained variant sites from the VCF files by using GATK SelectVariants^35–37^. We then filtered these variants by applying GATK Hard Filter (QD < 2.0, FS > 60.0, MQ < 40.0, MQRankSum < −12.5, ReadPosRankSum < −8.0). In addition, we only selected biallelic SNPs and removed any clustered SNPs defined by having 3 SNPs within 10bp.

### Filtering nucleotide sites

#### Filtering out the pseudoautosomal regions (PARs) of the X chromosome

Previous work showed that the PARs in canines span the first 6.59Mb of the X chromosome^40^. Therefore, we filtered out the PARs by removing any site that overlaps with the first 6.59Mb of the X chromosome. In humans, it was shown that genetic diversity does not drop abruptly at the PAR boundary^41^. Rather, genetic diversity decreases gradually over the PAR boundary and reaches nonPAR diversity past the PAR boundary^41^. One concern is that filtering out the PARs is not sufficient to avoid any inflation of X-linked variation. However, if this is the case, we would expect Q_π_ we calculated to be higher than the actual Q_π_. Therefore, Q_π_ less than 0.75 is not caused by not sufficiently filtering sites on the nonPARs.

#### Filtering sites that could be under the direct effect of selection

To control for the effects of direct selection, we removed sites that are potentially functional and therefore are more likely to be affected by purifying or positive selection. Specifically, we removed sites that overlap with a gene transcript as defined by Ensembl (gene transcripts include both exons and introns). We also removed sites that are conserved across species. To obtain conserved sites, we downloaded phastConsElements100way for hg19 from the UCSC Genome Browser and used liftOver command line tool to convert hg19 coordinates to canFam3 coordinates.

#### Filtering out sites that could be affected by linked selection

To control for the effect of natural selection on linked neutral sites, we employed a filtering criterion to remove sites near genes as defined by genetic distance to the nearest genes. We used the genetic distance map based on patterns of linkage disequilibrium from Auton et al. (2013) because this genetic map includes information for the X chromosome whereas the pedigree map from Campbell et al. (2016) does not have information on the X chromosome^42,43^. For each site that is outside of genes and conserved regions, we found its nearest gene in terms of physical distance. We then converted physical distance to genetic distance using the genetic map from Auton et al. (2013)^42^. Since we did not know *a priori* what the minimum genetic distance is required to remove sites near genes to control for linked selection, we used multiple thresholds. Specifically, we removed sites whose genetic distance to the nearest gene is less than 0.2 cM, less than 0.4 cM, less than 0.6 cM, less than 0.8 cM, and less than 1 cM.

#### Identifying sites that are alignable between dog and cat

Since we controlled for mutation rate variation by normalizing the uncorrected genetic diversity by dog-cat divergence, we identified regions of the genome that are alignable between dog and cat. We downloaded the pairwise alignment between dog and cat from the UCSC Genome browser^44^. We then generated BED files whose coordinates represent regions of the genome that are alignable between dog and cat. In summary, for our empirical analyses, we used regions of the genome that are (1) not affected directly by selection, (2) not affected by linked selection using multiple thresholds, (3) high in quality (see the section on filtering to obtain high quality sites above), and (4) alignable between dog and cat.

### Computing Q_π_

#### Computing uncorrected average pairwise differences between sequences (π)

We computed genetic diversity, *π*, defined as the average number of differences between pairs of sequences^45^: 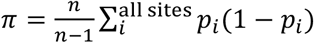where *p_i_* is the allele frequency and *n* is the number of alleles. For each region of the genome that satisfies the filtering criteria above, we computed π for the X chromosome and autosomes. To obtain the mean in diversity, π/site, we calculated: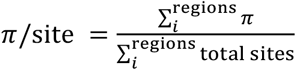

#### Computing dog-cat divergence

For each region of the genome that satisfies the filtering criteria above, we tabulated the number of DNA differences between dog and cat. To obtain the mean in divergence, we calculated 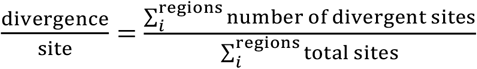

#### Computing male mutation bias

We computed male mutation bias (α) using divergence on the X chromosome and on the autosomes as follows: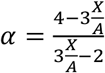

#### Computing corrected diversity

To control for variation in mutation rates across chromosomes, we normalized diversity by dog-cat divergence by dividing π/site by divergence/site.

#### Constructing 95% confidence interval by bootstrapping

We generated bootstrap replicates of the BED file that we used to compute genetic diversity and divergence by randomly selecting a fragment from the BED file with replacement. For each bootstrap replicate, the number of fragments chosen was equal to the number of fragments in the original BED file. We generated 1000 bootstrap replicates. For each of the 1000 bootstraps on the X chromosome, we computed uncorrected π, dog-cat divergence, and corrected π. We did the same calculations for each of the 1000 bootstraps on the autosomes. We then divided corrected π on the X chromosome by corrected π on the autosomes to obtain Q_π_. We calculated 95% confidence interval using 1000 bootstrapped values of π_X_, 1000 bootstrapped values of corrected π_A_, and 1000 bootstrapped values of corrected Q_π_ by selecting the values at the 2.5 and 97.5 percentiles.

### Computing Q_FST_

#### Computing F_ST_

We computed Weir and Cockerham’s F_ST_ for each pair of populations using the *SNPRelate* package implemented in *R*^47^. For dog-to-dog comparison, we computed F_ST_ for German Shepherds and Tibetan Mastiffs, German Shepherds and Pooled Breed Dogs, and Tibetan Mastiff and Pooled Breed Dogs. For wolf-to-wolf comparison, we computed F_ST_ for Arctic Wolves and Grey Wolves. Since the number of individuals differs between populations, we subsampled such that there were four individuals in each population (Supplementary Table 8). We computed F_ST_ for the X chromosome and for the autosomes.

#### Computing Q_FST_

We computed Q_FST_ using: 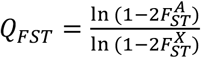

#### Constructing 95% confidence interval by bootstrapping

Since the input to *SNPRelate* to calculate F_ST_ is a VCF file format, we generated 1000 bootstrapped VCF files by randomly selecting variants from the VCF file with replacement. The number of variants selected for each bootstrapped VCF is equal to the number of variants in the empirical VCF file. For each bootstrapped VCF, we computed F_ST_ and Q_FST_ as explained above. From the 1000 values of bootstrapped Q_FST_, we then calculated 95% confidence interval by selecting the values at the 2.5 and 97.5 percentiles.

### Modeling framework to estimate the N_X_ / N_A_ ratio (C)

#### Obtaining the site frequency spectrum (SFS)

We computed the folded SFSs using Equation 1.2 of Wakely’s An Introduction to Coalescent Theory^48^, reproduced as follows:

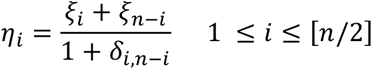

where ξ_*i*_ is the number of sites where the alternate allele is present at *i* copies, *δ,_n-i_* is equal to 0 when *i ≠ n − i* and is equal to 1 when *i = n − i*. For each population and for each threshold to remove linked neutral sites (>0.4 cM, >0.6 cM, >0.8 cM, and >1 cM), we computed the folded SFSs for the X chromosome and autosomes.

#### Computing mutation rates

We utilized dog-cat divergence to infer the mutation rates for the X chromosome and autosomes. Specifically, 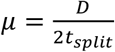, where D is the divergence/site between dog and cat (see Computing dog-cat divergence section above) and *t_split_* is the split time between dog and cat in unit of generation. We used 54 million years as the split time between dog and cat and a generation time of 3 years per generation^49,50^.

The estimates of mutation rates are in the same order of magnitude as estimate from ancient DNA (Supplementary Table 4)^8^.

#### Inferring demographic parameters

We inferred demographic parameters from the autosomal data (SFSs on the autosomes) using a maximum likelihood framework as implemented in *fastsimcoal2*^33^. We specified a bottleneck demographic model and inferred four parameters: N_ANC_ which is the population size in the ancestral population, N_BOT_ which is the population size during the bottleneck, N_CUR_ which is the population size in the current day, and T_BOT_ which is the duration between the end of the bottleneck and current day (Supplementary Figure 1). Further, we repeated the inference of the previous four parameters for values BOT_DUR_ (the duration of the bottleneck) ranging from 75 to 100 generations (Supplementary Figure 1) and chose the value that yielded the highest likelihood. We implemented this procedure for each population and for thresholds of >0.4 cM, >0.6 cM, >0.8 cM, and >1 cM to remove linked sites. The demographic parameters that maximized the likelihood are summarized in Supplementary Table 5.

#### Inferring N_X_/N_A_ ratio (C)

To account for differences in population size between the X chromosome and autosomes, we scaled the population size on the X chromosome to that on the autosomes by a constant factor we called C, where N_X_ = CN_A_. To find the maximum likelihood estimate of C, we searched over a grid for values of C, including 0.75, to find a value that resulted in the highest likelihood. Because the number of SNPs at particular frequencies contains substantial information about demography, we used a Poisson likelihood for the number of SNPs in each entry of the SFS to compute the Poisson log-likelihood as in Beichman et al. (2017)^51^.

#### Accessing fit of MLEs of C to π

We computed diversity from the simulated SFSs under the demographic models fit to the autosomes using the MLEs of *C* (Table 1) and compared that to the empirical uncorrected diversity.

## Data availability

All scripts can be found at https://github.com/tnphung/SexBiased. SRA numbers for *fastq* files are listed in Supplementary Table 1. Post base quality score calibration (BQSR) BAM files and VCF files are available upon request.

## ACKNOWLEDGEMENTS

We thank Jacqueline Robinson for providing the sequencing data for the Arctic Wolves and Christian Huber for helpful discussions. This work was supported by the National Institute of General Medical Sciences (NIGMS) of the National Institutes of Health (NIH) grant R35GM119856 to K.E.L and NIGMS grant R35GM124827 to M.A.W.S. T.N.P. was supported by NIH-NCI National Cancer Institute T32CA201160.

## REFERENCES

1. Hemmer, H. Domestication: The Decline of Environmental Appreciation. (Cambridge University Press, 1990).

2. Freedman, A. H. et al. Genome sequencing highlights the dynamic early history of dogs. PLoS Genet. 10, e1004016 (2014).

3. Freedman, A. H., Lohmueller, K. E. & Wayne, R. K. Evolutionary history, selective sweeps, and deleterious variation in the dog. Annu. Rev. Ecol. Evol. Syst. 47, 73–96 (2016).

4. Ostrander, E. A., Wayne, R. K., Freedman, A. H. & Davis, B. W. Demographic history, selection and functional diversity of the canine genome. Nat. Rev. Genet. 18, 705–720 (2017).

5. Freedman, A. H. & Wayne, R. K. Deciphering the origin of dogs: from fossils to genomes. Annu. Rev. Anim. Biosci. 5, 281–307 (2017).

6. Boyko, A. R. The domestic dog: man’s best friend in the genomic era. Genome Biol. 12, 216 (2011).

7. vonHoldt, B. M. et al. A genome-wide perspective on the evolutionary history of enigmatic wolf-like canids. Genome Res. 21, 1294–1305 (2011).

8. Larson, G. et al. Rethinking dog domestication by integrating genetics, archeology, and biogeography. Proc. Natl. Acad. Sci. 109, 8878–8883 (2012).

9. Thalmann, O. et al. Complete mitochondrial genomes of ancient canids suggest a European origin of domestic dogs. Science 342, 871–874 (2013).

10. Botigué, L. R. et al. Ancient European dog genomes reveal continuity since the Early Neolithic. Nat. Commun. 8, 16082 (2017).

11. Frantz, L. A. F. et al. Genomic and archaeological evidence suggest a dual origin of domestic dogs. Science 352, 1228–1231 (2016).

12. Drake, A. G., Coquerelle, M. & Colombeau, G. 3D morphometric analysis of fossil canid skulls contradicts the suggested domestication of dogs during the late Paleolithic. Sci. Rep. 5, 8299 (2015).

13. Wilson Sayres, M.A. Genetic diversity on the sex chromosomes. Genome Biol. Evol. 10, 1064– 1078 (2018).

14. Vonholdt, B. M. et al. The genealogy and genetic viability of reintroduced Yellowstone grey wolves. Mol. Ecol. 17, 252–274 (2008).

15. Casanova wolves | Natural History. Available at: https://retrieverman.net/2010/12/17/casanova-wolves/. (Accessed: 26th June 2018)

16. Baker, P. J., Funk, S. M., Bruford, M. W. & Harris, S. Polygynandry in a red fox population: implications for the evolution of group living in canids? Behav. Ecol. 15, 766–778 (2004).

17. Sillero-Zubiri, C., Gottelli, D. & Macdonald, D. W. Male philopatry, extra-pack copulations and inbreeding avoidance in Ethiopian wolves (*Canis simensis*). Behav. Ecol. Sociobiol. 38, 331–340 (1996).

18. Sundqvist, A.-K. et al. Unequal contribution of sexes in the origin of dog breeds. Genetics 172, 1121–1128 (2006).

19. Ostrander, E. A. & Kruglyak, L. Unleashing the canine genome. Genome Res. 10, 1271–1274 (2000).

20. Emery, L. S., Felsenstein, J. & Akey, J. M. Estimators of the human effective sex ratio detect sex biases on different timescales. Am. J. Hum. Genet. 87, 848–856 (2010).

21. Webster, T. H. & Wilson Sayres, M. A. Genomic signatures of sex-biased demography: progress and prospects. Curr. Opin. Genet. Dev. 41, 62–71 (2016).

22. Hammer, M. F., Mendez, F. L., Cox, M. P., Woerner, A. E. & Wall, J. D. Sex-biased evolutionary forces shape genomic patterns of human diversity. PLoS Genet. 4, e1000202 (2008).

23. Keinan, A., Mullikin, J. C., Patterson, N. & Reich, D. Accelerated genetic drift on chromosome X during the human dispersal out of Africa. Nat. Genet. 41, 66–70 (2009).

24. Hammer, M. F. et al. The ratio of human X chromosome to autosome diversity is positively correlated with genetic distance from genes. Nat. Genet. 42, 830–831 (2010).

25. Arbiza, L., Gottipati, S., Siepel, A. & Keinan, A. Contrasting X-linked and autosomal diversity across 14 human populations. Am. J. Hum. Genet. 94, 827–844 (2014).

26. Gou, X. et al. Whole-genome sequencing of six dog breeds from continuous altitudes reveals adaptation to high-altitude hypoxia. Genome Res. 24, 1308–1315 (2014).

27. Marsden, C. D. et al. Bottlenecks and selective sweeps during domestication have increased deleterious genetic variation in dogs. Proc. Natl. Acad. Sci. 113, 152–157 (2016).

28. Li, W. H., Yi, S. & Makova, K. Male-driven evolution. Curr Opin Genet Dev 12, (2002).

29. Wilson Sayres, M. A. & Makova, K. D. Genome analyses substantiate male mutation bias in many species. BioEssays News Rev. Mol. Cell. Dev. Biol. 33, 938–945 (2011).

30. Lindblad-Toh, K. et al. Genome sequence, comparative analysis and haplotype structure of the domestic dog. Nature 438, 803–819 (2005).

31. Narang, P., Wilson Sayres, M. A. Variable autosomal and X divergence near and far from genes affects estimates of male mutation bias in great apes. Genome Biol. Evol. 8, 3393–3405 (2016).

32. Pool, J. E. & Nielsen, R. Population size changes reshape genomic patterns of diversity. Evolution 61, 3001–3006 (2007).

33. Excoffier, L., Dupanloup, I., Huerta-Sánchez, E., Sousa, V. C. & Foll, M. Robust demographic inference from genomic and SNP data. PLoS Genet 9, e1003905 (2013).

34. Stahler, D. R., MacNulty, D. R., Wayne, R. K., vonHoldt, B. & Smith, D. W. The adaptive value of morphological, behavioural and life-history traits in reproductive female wolves. J. Anim. Ecol. 82, 222–234

35. McKenna, A. et al. The Genome Analysis Toolkit: A MapReduce framework for analyzing next-generation DNA sequencing data. Genome Res. 20, 1297–1303 (2010).

36. DePristo, M. A. et al. A framework for variation discovery and genotyping using next-generation DNA sequencing data. Nat. Genet. 43, 491–498 (2011).

37. Van der Auwera, G. A. et al. From FastQ data to high confidence variant calls: the Genome Analysis Toolkit best practices pipeline. Curr. Protoc. Bioinforma. 43, 11.10.1-33 (2013).

38. Picard Tools - By Broad Institute. Available at: http://broadinstitute.github.io/picard/. (Accessed: 9th March 2018)

39. Li, H. Aligning sequence reads, clone sequences and assembly contigs with BWA-MEM. ArXiv13033997 Q-Bio (2013).

40. Young, A. C., Kirkness, E. F. & Breen, M. Tackling the characterization of canine chromosomal breakpoints with an integrated in-situ/in-silico approach: The canine PAR and PAB. Chromosome Res. 16, 1193–1202 (2008).

41. Cotter, D. J., Brotman, S. M. & Wilson Sayres, M. A. Genetic diversity on the human X chromosome does not support a strict pseudoautosomal boundary. Genetics 203, 485–492 (2016).

42. Auton, A. et al. Genetic recombination is targeted towards gene promoter regions in dogs. PLoS Genet 9, e1003984 (2013).

43. Campbell, C. L., Bhérer, C., Morrow, B. E., Boyko, A. R. & Auton, A. A pedigree-based map of recombination in the domestic dog genome. G3 GenesGenomesGenetics 6, 3517–3524 (2016).

44. Kent, W. J. et al. The Human Genome Browser at UCSC. Genome Res. 12, 996–1006 (2002).

45. Tajima, F. Evolutionary relationship of DNA sequences in finite populations. Genetics 105, 437– 460 (1983).

46. Link, V., Aguilar-Gómez, D., Ramírez-Suástegui, C., Hurst, L. D. & Cortez, D. Male mutation bias is the main force shaping chromosomal substitution rates in monotreme mammals. Genome Biol. Evol. 9, 2198–2210 (2017).

47. Zheng, X. et al. A high-performance computing toolset for relatedness and principal component analysis of SNP data. Bioinforma. Oxf. Engl. 28, 3326–3328 (2012).

48. Wakely, J. Coalescent Theory: An Introduction. (Macmillan Learning, 2016).

49. Hedges, S. B., Marin, J., Suleski, M., Paymer, M. & Kumar, S. Tree of life reveals clock-like speciation and diversification. Mol. Biol. Evol. 32, 835–845 (2015).

50. Hedges, S. B., Dudley, J. & Kumar, S. TimeTree: a public knowledge-base of divergence times among organisms. Bioinforma. Oxf. Engl. 22, 2971–2972 (2006).

51. Beichman, A. C., Phung, T. N. & Lohmueller, K. E. Comparison of single genome and allele frequency data reveals discordant demographic histories. G3 Genes Genomes Genet. 7, 3605–3620 (2017).

